# Evolving alterations of structural organization and functional connectivity in feedforward neural networks after induced P301L tau mutation

**DOI:** 10.1101/2023.09.12.557339

**Authors:** Janelle S. Weir, Katrine Sjaastad Hanssen, Nicolai Winter-Hjelm, Axel Sandvig, Ioanna Sandvig

**Affiliations:** Department of Neuromedicine and Movement Science, Faculty of Medicine and Health Sciences, Norwegian University of Science and Technology (NTNU), Trondheim, Norway; Department of Community Medicine and Rehabilitation, Umeå University, Umeå, Sweden; Department of Neurorehabilitation, Umeå University Hospital, Umeå, Sweden

**Keywords:** Cortical network, neural engineering, electrophysiology, mutated tau, self-organization, microfluidic chip

## Abstract

Reciprocal structure–function relationships underlie both healthy and pathological behaviors in complex neural networks. Thus, understanding neuropathology and network dysfunction requires a thorough investigation of the complex interactions between structural and functional network reconfigurations in response to perturbation. Such adaptations are often difficult to study *in vivo*. For example, subtle, evolving changes in synaptic connectivity, transmission, and the electrophysiological shift from healthy to pathological states, as for example alterations that may be associated with evolving neurodegenerative disease, such as Alzheimeŕs, are difficult to study in the brain. Engineered *in vitro* neural networks are powerful models that enable selective targeting, manipulation, and monitoring of dynamic neural network behavior at the micro- and mesoscale in physiological and pathological conditions. In this study, we engineered feedforward cortical neural networks using two-nodal microfluidic devices with controllable connectivity interfaced with microelectrode arrays (mMEAs). We induced P301L mutated tau protein to the presynaptic node of these networks and monitored network dynamics over three weeks. Induced perturbation resulted in altered structural organization and extensive axonal retraction starting in the perturbed node. Perturbed networks also exhibited functional changes in intranodal activity, which manifested as an overall decline in both firing rate and bursting activity, with a progressive increase in synchrony over time, and decrease in internodal signal propagation between pre- and postsynaptic nodes. These results provide insights into dynamic structural and functional reconfigurations at the micro- and mesoscale as a result of evolving pathology and illustrate the utility of engineered networks as models of network function and dysfunction.

## 1. Introduction

One of the main pathological hallmarks of Alzheimeŕs disease (AD) is the accumulation of neurofibrillary tangles in the brain resulting from hyperphosphorylation of the microtubule associated protein tau (Braak et al., 1991). Tau proteins are mainly found in the axons of neurons (Götz et al., 2019) where they stabilize microtubules and promote their assembly (Baas et al., 2019; Qiang et al., 2018). However, under pathological conditions, tau becomes hyperphosphorylated, causing widespread morphological axonal disruption (Jackson et al., 2017; Kopeikina et al., 2013), ultimately affecting synaptic transmission and network function. Evolving neuropathology can also have widespread implications on the structural properties of neural networks, and drive changes in the functional interactions among network components. The high interconnectivity of the brain implies that neuronal axons and synapses may act as conduits for the spread of such pathology from affected areas, causing progressive, widespread disruption to the structural and functional integrity of the network (Adams et al., 2019; Kuhl, 2019). It has also been shown that hyperphosphorylated tau triggers hypoactivity in neurons (Ittner et al., 2011; Menkes-Caspi et al., 2015; Polanco et al., 2018; Wang et al., 2017), thus disturbing excitatory-inhibitory balance, and disrupting synaptic transmission and global integration across the network.

To elucidate how relevant pathological mechanisms gradually affect the structure and function of neural networks, it is imperative to study the underlying processes at the micro- and mesoscale. Such investigations are highly challenging, or de facto not feasible *in vivo* due to the size and complexity of the brain. This challenge can be overcome with the application of advanced cellular models such as engineered neural networks. Such networks develop with progressively increasing structural and functional complexity over time, recapitulating fundamental aspects of neural network behavior as seen in the brain (Collingridge et al., 2010; Valderhaug et al., 2021; van de Wijdeven et al., 2019; Winter-Hjelm N, 2023). Engineered *in vitro* models thus enable longitudinal studies of dynamic network behavior and allow for selective perturbation and monitoring of network responses at the micro- and mesoscale level (Bauer et al., 2022; Bruno et al., 2020; Fiskum et al., 2021; Gribaudo et al., 2019; Nonaka et al., 2011; Valderhaug et al., 2021; Valderhaug et al., 2024; Weir et al., 2023).

In the present study, we longitudinally investigated structural and functional changes in engineered two-nodal feedforward cortical neural networks following induced expression of human mutated tau in the presynaptic nodes. Our primary aim was to monitor and identify dynamic changes in neurite organization and electrophysiological activity of networks with induced perturbation and compare this with unperturbed control networks. The two-nodal feedforward configuration of the engineered network enabled us to observe structural and functional reconfigurations in response to the perturbation in the affected node, while simultaneously monitoring dynamic changes in the postsynaptic node. Our study provides significant insights into neural network dynamics, demonstrating how networks adapt and reconfigure in response to human mutated tau perturbations, relevant to neurodegenerative diseases like Alzheimer’s. It shows that mutated tau expression disrupts internodal connectivity, causing progressive neurite retraction and reduced neural activity, while increasing network synchrony, a potential marker of pathological states. The absence of these changes in control networks confirms the perturbation’s impact, highlighting the importance of neurite integrity for network function. Additionally, our study shows that the use of engineered neural networks as models proves valuable for studying dynamic micro-mesoscale network reconfigurations of neural dysfunction, emphasizing their role in investigating mechanisms of neural resilience and vulnerability.

## 2. Methods

### 2.1. *In vitro* neural networks

An experimental timeline can be found in **Figure 1**. For this study, we used in-house developed microfluidic chips interfaced with microelectrode arrays (mMEAs; n=7) and 8-well chambered slides (Ibidi, 80841; n=2). Design and fabrication of the mMEAs was conducted as reported previously by our group (Winter-Hjelm N, 2023). Briefly, two compartments (from here on referred to as nodes) (5 mm wide/60 µm high) were connected by 20 microtunnels (700 µm long/10 µm wide/5 µm high) designed to promote unidirectional axonal outgrowth from the presynaptic to the postsynaptic node. Tesla valve microtopographies were included in the microtunnels to redirect axons from the postsynaptic node back to their chamber of origin. Furthermore, axon traps were included on the postsynaptic side to misguide outgrowing axons and prevent them from entering the microtunnels. To prevent neuronal somata from entering the microtunnels, 4 µm pillars with 4 µm interspacing were positioned within the presynaptic node. Impedance measurements and sterilization of the mMEAs were conducted as reported previously (Winter-Hjelm N, 2023). Prior to plating of the cells, nodes were coated with 0.1 mg/mL Poly-l-Ornithine (PLO; Sigma-Aldrich, #P4957) for 30 min, subsequently replenished with fresh PLO for another 2 h at 37°C/5% CO2. Following this, all PLO was discarded, and the surfaces rinsed three times with distilled Milli-Q-water. Subsequently, the platforms were coated with laminin solution consisting of 16 µg/mL Mouse Natural Laminin (Gibco, #23017015) diluted in PBS (Sigma-Aldrich, D8537) for first 30 min, before being replenished with fresh laminin solution for another 2 h at 37°C/5% CO2. To ensure proper flow of coating solution through the microtunnels, a hydrostatic pressure gradient was applied during all coating steps. Laminin solution was discarded and replaced by astrocyte media consisting of DMEM (Gibco™, 11885084) supplemented with 15% Fetal Bovine Serum (Sigma-Aldrich, F9665) and 2% Penicillin-Streptomycin (Sigma-Aldrich, #P4333) for 10 min at 37°C/5% CO_2_ before rat primary cortical astrocytes (Gibco, #N7745100) were plated at a density of 100 cells/mm^2^ 48 hours prior to plating of neurons. Subsequently, rat primary cortical neurons (Gibco, #A1084001) were plated at a density of 1,000 cells/mm^2^ in neuronal media consisting of Neurobasal Plus Medium (Gibco™, A3582801) supplemented with 2% B27 Plus Medium (Gibco™, A358201), 2.5 mL/L Gluta-Max (Gibco™, 35050038) and 1% Penicillin-Streptomycin (Sigma-Aldrich, P4333). 4 and 24 hours after plating the neurons, 90% of the media was replenished. Thereafter, 50% of the media was replenished every 2 days.

**Figure 1.**
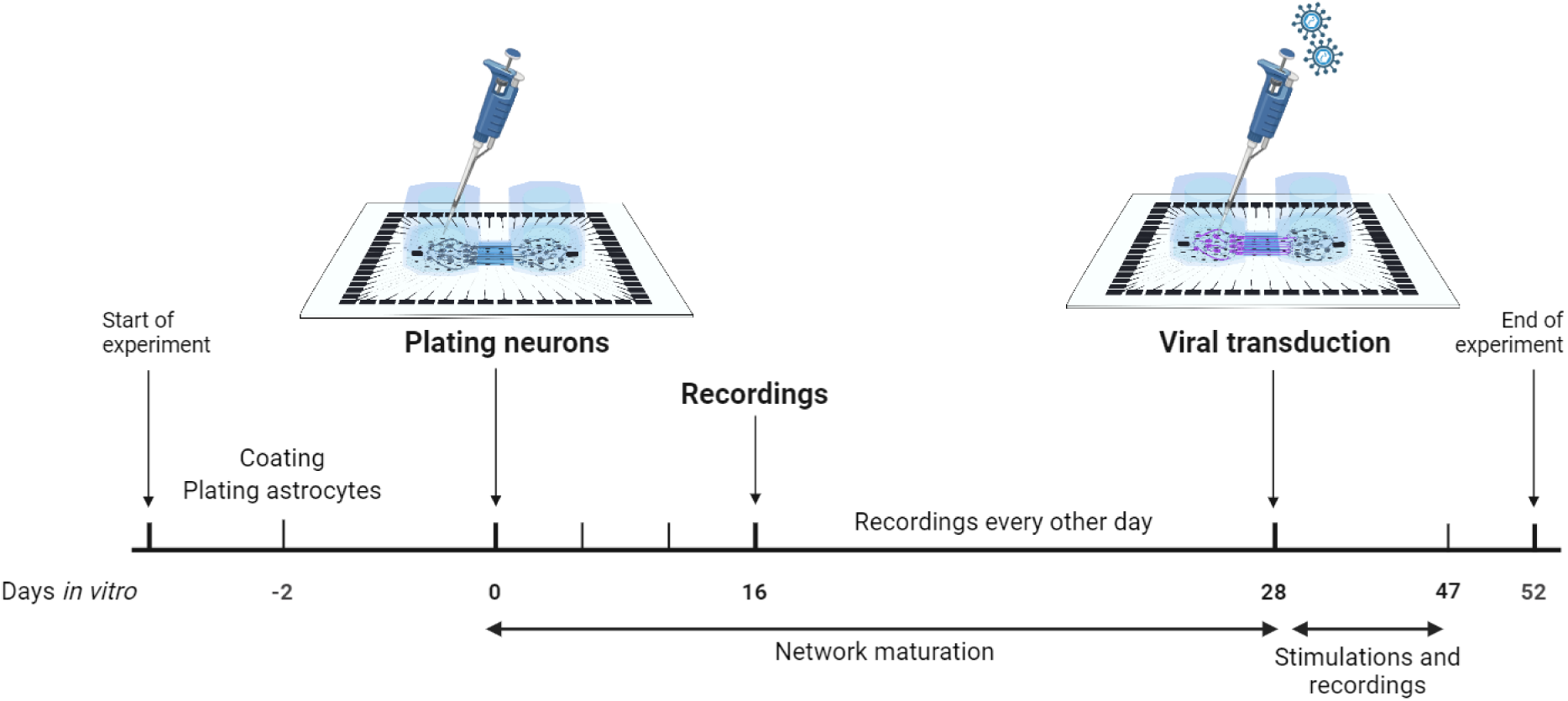
Schematic of the *in vitro* experimental timeline for establishing networks, transduction, and electrophysiological recordings. *Created with BioRender.com*.

#### 2.1.1. AAV8 – GFP – 2A P301L Tau production and *in vitro* transduction

Adeno associated virus (AAV) vector production and purification was performed in-house at the Viral Vector Core Facility (Kavli Institute for Systems Neuroscience, NTNU). Titering of the viral stock was determined as approximately 10^11^ vg/mL. The viral stock was divided into 20 µl aliquots and stored at -80°C . Aliquots designated for use were thawed on ice, while any remaining virus was aliquoted and stored at -80°C for future use. Each aliquot was thawed a maximum of 3 times. Viral units were introduced to healthy networks at 28 DIV to induce perturbation. Neurons were transduced by removing 70% of the cell media from the presynaptic nodes and directly adding 3 x 10^2^ viral units per neuron diluted in cell media. After addition of the virus, the cultures were gently agitated for 30 s to ensure proper distribution and then incubated for 4 h. Following our previously published *in-vitro* AAV vector study (Weir et al., 2023), we evaluated varying viral concentrations ranging from 1 x 10^2^ to 5 x 10^3^ viral units per neurons. The final titer of viral units per neuron was decided by selecting the lowest concentration that achieved over 50% transduced cells with networks surviving more than 3 weeks following transduction. Afterwards, each chamber was topped up with fresh media without antibiotics and incubated for an additional 20 h at 37°C/5% CO2. After the incubation period, 50% changes of the media were carried out every second day. To ensure comparable conditions with the control networks, a 70% media change was also conducted in the control unperturbed networks at the same time as the addition of AAV P301L to the perturbed networks. The viral vector encoded a GFP fluorescent tag for easy visualization of transduction efficiency.

### 2.2. Immunocytochemistry

To assess network maturity including the expression of structural neuronal markers such as microtubule associated protein, immunocytochemistry (ICC) was performed as follows. Prior to immunostaining, cell media were aspirated and discarded from the culture plates, and the cultures were rinsed once with Dulbecco’s Phosphate Buffered Saline (DPBS; Thermo Fischer Scientific, Cat #14040-117). Following this, networks were fixed with 4% Paraformaldehyde (Sigma-Aldrich, #P6148) for 10 min at room temperature followed by 3x10 min washes with DPBS. Further, all DPBS were discarded and replaced by blocking solution consisting of 5% Goat serum (Sigma-Aldrich, Cat# G9023) and 0.3% Triton-X (Thermo Fischer Scientific, Cat# 85111) in DPBS. Next, primary antibodies at the indicated concentrations (**Table 1**) were added in a buffer of 0.01% Triton-X and 1% Goat Serum in DPBS overnight at 4°C. The following day, primary antibody solution was discarded, and cultures were rinsed 3x5 min with DBPS before secondary antibodies at the indicated concentrations (**Table 1**) were added in a buffer of 0.01% Triton-X and 1% Goat Serum in DPBS for 2 h at room temperature. For staining cellular nuclei, Hoechst dye (bisbenzimide H 33342 trihydrochloride; Sigma-Aldrich, Cat# 14533) was added at 1:10.000 dilution for the last 10 min of the secondary antibody incubation. Samples from the 8-well Ibidi chips were washed with PBS, mounted on glass cover slides with anti-fade fluorescence mounting medium (Abcam, Cat#Ab104135) whereas microfluidic chips were filled with distilled Milli-Q-water before imaging.

**Table 1.** List of primary and secondary antibodies with concentrations.

| Primary antibodies | Concentration | Supplier |
| --- | --- | --- |
| Ck Map2 | 1:1000 | Abcam, #Ab5392 |
| Rb Phospho-Tau (Thr217) (pTau) | 1:1000 | Invitrogen, #44-744 |
| Ms AT8 (Ser202/205) | 1:1000 | Invitrogen, #MN1020 |
| Rb GAD65/67 | 1:100 | Abcam, #Ab183999 |
| Ms Calmodulin (CaMKIIa) | 1:200 | Invitrogen, #MA3-918 |
| Rb GFAP | 1:500 | Abcam, #Ab278054 |
| Secondary antibodies | Concentration | Supplier |
| Goat-Anti-Mouse Alexa Fluor 674 | 1:1000 | Invitrogen, #A21236 |
| Goat-Anti-Rabbit Alexa Fluor 488 | 1:1000 | Invitrogen, #A21244 |
| Goat-Anti-Chicken Alexa Fluor 568 | 1:1000 | Abcam, #Ab175477 |

*Ms: Mouse, Rb: rabbit, Ck: chicken.*

#### 2.2.1 Imaging

All samples from ICC were imaged using an EVOS M5000 microscope (Thermo Fischer Scientific, #AMF5000) connected to a LED light source and using an Olympus 20x/0.75 NA objective (N1480500), with the following filter sets/channels: DAPI (AMEP4650), CY5 (AMEP4656), GFP (AMEP4651) and TxRed (AMEP4655). Phase contrast images were taken using a Zeiss Axio Vert V.1 brightfield 20x/53 WD/0.4 NA objective with an Axiocam 202 mono. Images were processed using Adobe Photoshop 2020. To quantify for fluorescent intensity profile of AT8 and pTau in cell body clusters and axonal bundles in both perturbed and control unperturbed networks, fluorescent signal was measured in Fiji/ImageJ2. A total of six clusters of cell bodies and five axonal bundles were selected as regions of interest and assessment was done for total fluorescent signal (mean intensity/pixel) measured by mean grayscale values. Statistical analysis of the mean grayscale values was done using Statistical Package for the Social Sciences (SPSS) version 29.0.0.0.

### 2.3. Electrophysiological recordings

Electrophysiological activity of neural networks on mMEAs (n=7) was recorded using a MEA2100 recording system (Multichannel Systems, MCS, Reutlingen, Germany) at a sampling rate of 25,000 Hz. A 3D-printed in-house made plastic cap covered by a gas-permeable membrane was used to keep the cultures sterile during recordings. The stage temperature was set to 37°C (TC01, Multichannel Systems) and the cultures were allowed to equalize on the stage for 5 min before spontaneous electrophysiological activity was recorded for 15 min. All networks were recorded 24 h after media changes on the following days *in vitro* (DIV): 16, 20, 24, 26, 28, 31, 33, 35, 37, 39, 41, 43, 45 and 47. From 28 DIV onwards, networks were electrically stimulated and simultaneously recorded for 1 minute (directly following the 15 min recordings of spontaneous electrophysiological activity). Electrical stimulations were applied to one presynaptic electrode within each of the mMEAs. The electrode with the highest detected mean firing rate (Pasquale et al., 2010) during the 15 min recording at 16 DIV for each of the mMEAs was selected for stimulations. Stimulations consisted of a train of 30 spikes at ± 600 mV (positive phase first) of 200 μs duration with an interspike interval of 5 s. This was done according to previous studies demonstrating that persistent electrical stimulation in *in vitro* networks can increase network activity over time, resulting in enhanced evoked action potentials and an increased frequency of spikes in bursts (Brewer et al., 2009; Ide et al., 2010). Raw data was converted to an .h5 Hierarchical Data Format using Multichannel DataManager (V.1.14.4.22018) system and imported to Matlab R2021b for further analyses using adapted and custom-made scripts.

### 2.4. Data analysis

Electrophysiology data analysis was done as previously described (Winter-Hjelm N, 2023). Briefly, raw data was filtered using a 4^th^ order Butterworth bandpass filter removing low frequency fluctuations below 300 Hz and high frequency noise above 3000 Hz. A notch filter was used to remove 50 Hz noise caused by the power supply mains. Stimulation data was filtered using the SALPA filter (Wagenaar et al., 2002), and each stimulation time point was followed by 15 ms blanking. Spike detection was conducted using the Precise Timing Spike Detection (PTSD) algorithm (Maccione et al., 2009) with a threshold of 11 times the standard deviation, a maximum peak duration set to 1 ms and a refractory period of 1.6 ms. Burst detection was conducted using the logISI approach (Pasquale et al., 2010), with a minimum of four consecutive spikes set as a minimum for a burst to be detected, and a hard threshold of 100 ms. Network bursts were detected using the logIBEI approach (Pasquale et al., 2010), with at least 20% of active electrodes required to participate during the span of a network burst. Network synchrony was measured using the coherence index (Timme et al., 2018).

### 2.5 Statistical analysis

Statistical Package for the Social Sciences (SPSS) version 29.0.0.0 was utilized for all statistical analyses. For comparison of the repeated measures, we used Generalized Linear Mixed-Effect Models (GLMMs) with network type (i.e., controls versus perturbed networks) as a fixed effect, and the network characteristics as targets. The network age (DIV) was used as a random effect. Only data from 31 DIV onwards were included in the analysis to specifically compare changes in network characteristics following perturbation to the unperturbed control networks. A gamma probability distribution with a log link function was chosen as the linear model. This selection was based on the Akaike information criterion and initial assessment of distribution fit to the predicted values. For multiple comparisons, we used sequential Bonferroni adjustment.

## 3. Results

### 3.1. Engineered feedforward cortical networks show intra- and internodal connectivity

Prior to perturbation, we validated that neurons were structurally connected with each other within and across the nodes and expressed markers for excitatory and inhibitory synaptic transmission. Immunocytochemistry performed at 22 DIV revealed expression of the microtubule-associated protein 2 marker, MAP2, the inhibitory neuronal marker glutamic acid decarboxylase 65/67, (GAD65/67) (Fukuda et al., 1997) (**Figure 2A**), and the excitatory neuronal marker calcium-calmodulin (CaM) – dependent protein kinase II, (CaMKII) (Takao et al., 2005) (**Figure 2B**). The presence of these markers indicated that the networks consisted of mature neurons with the capacity for excitatory and inhibitory synaptic transmission, a crucial aspect for achieving structural and functional network maturity.

**Figure 2.**
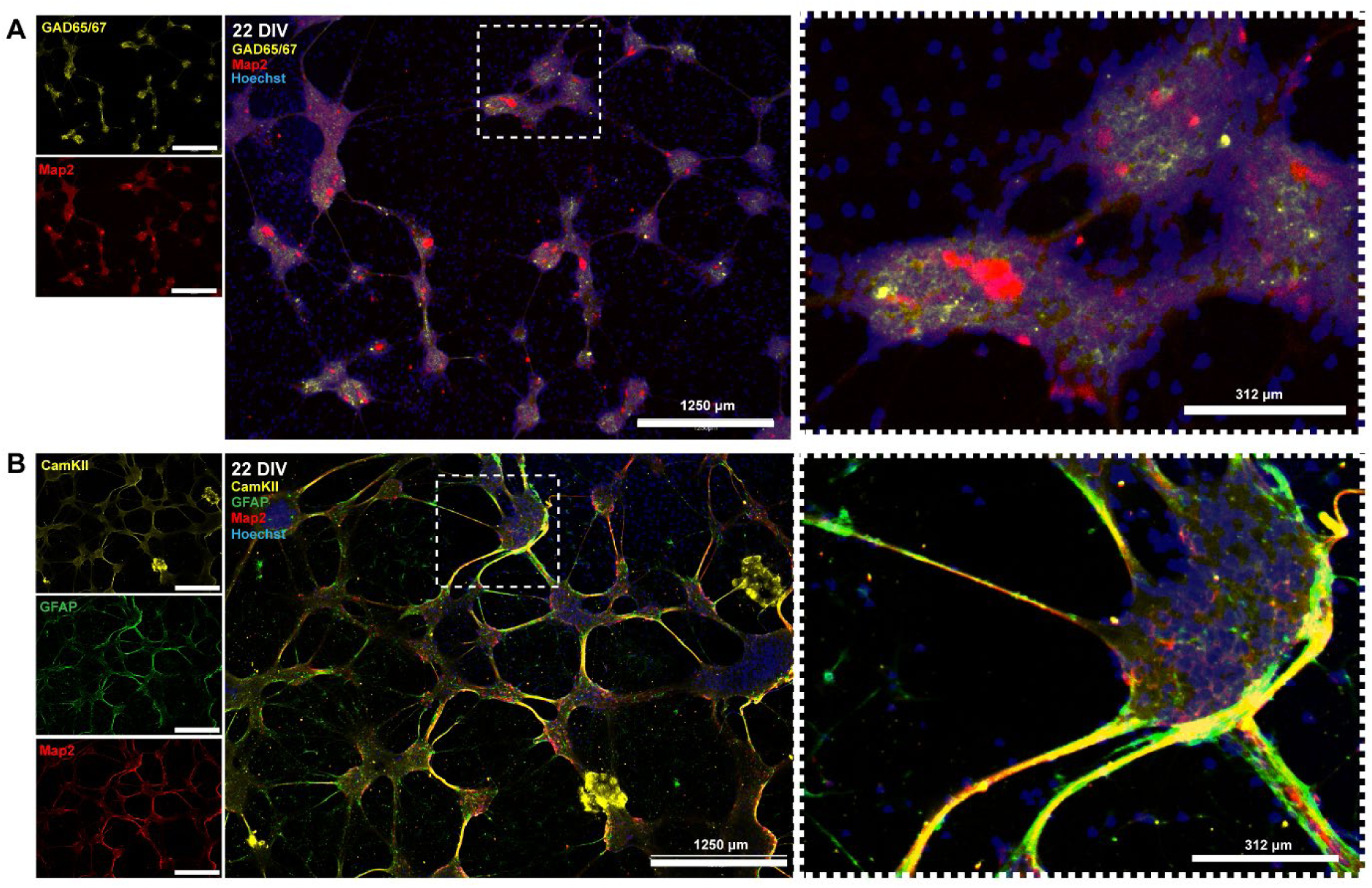
Developing neuronal mechanisms for excitatory and inhibitory synaptic transmission at 22 DIV. **(A)** Neural networks positively immunolabeled for the neuronal marker MAP2 and the inhibitory neuron marker GAD65/67, **(B)** and the excitatory neuron marker CaMKII. An antibody against Glial fibrillary acidic protein (GFAP) was used to label astrocytes in the network. DIV; days in vitro. Scale bar 1250 μm; (magnified area 312 μm).

Our results also showed that the neural networks organized into densely interconnected intranodal architectures, with outgrowing neurites in the microtunnels (**Figure 3A**). To investigate whether the engineered networks were functionally connected, we monitored and recorded electrophysiological activity commencing at 16 DIV. By 26 DIV, the neural networks displayed a mature electrophysiological profile in line with previous studies (Chiappalone et al., 2006; Winter-Hjelm N, 2023). Specifically, raster plots of spontaneous activity within the network revealed both isolated spikes and synchronized bursts (**Figure 3B**), while correlation matrices of network connectivity showed strong connectivity within pre- and postsynaptic nodes, as well as functional connectivity between the nodes (**Figure 3C**). To further validate internodal connectivity, we applied electrical stimulations to the electrode with the highest firing rate within the presynaptic node at 26 DIV. We recorded and assessed the evoked activity in the postsynaptic node, which revealed that electrical stimulations caused a spiking response in the presynaptic node, followed by an 40 ms (average) time delay before a subsequent spiking response was observed in the postsynaptic node (**Figure 3D**). This effectively demonstrated that functional connectivity was established between the-pre and postsynaptic nodes.

**Figure 3.**
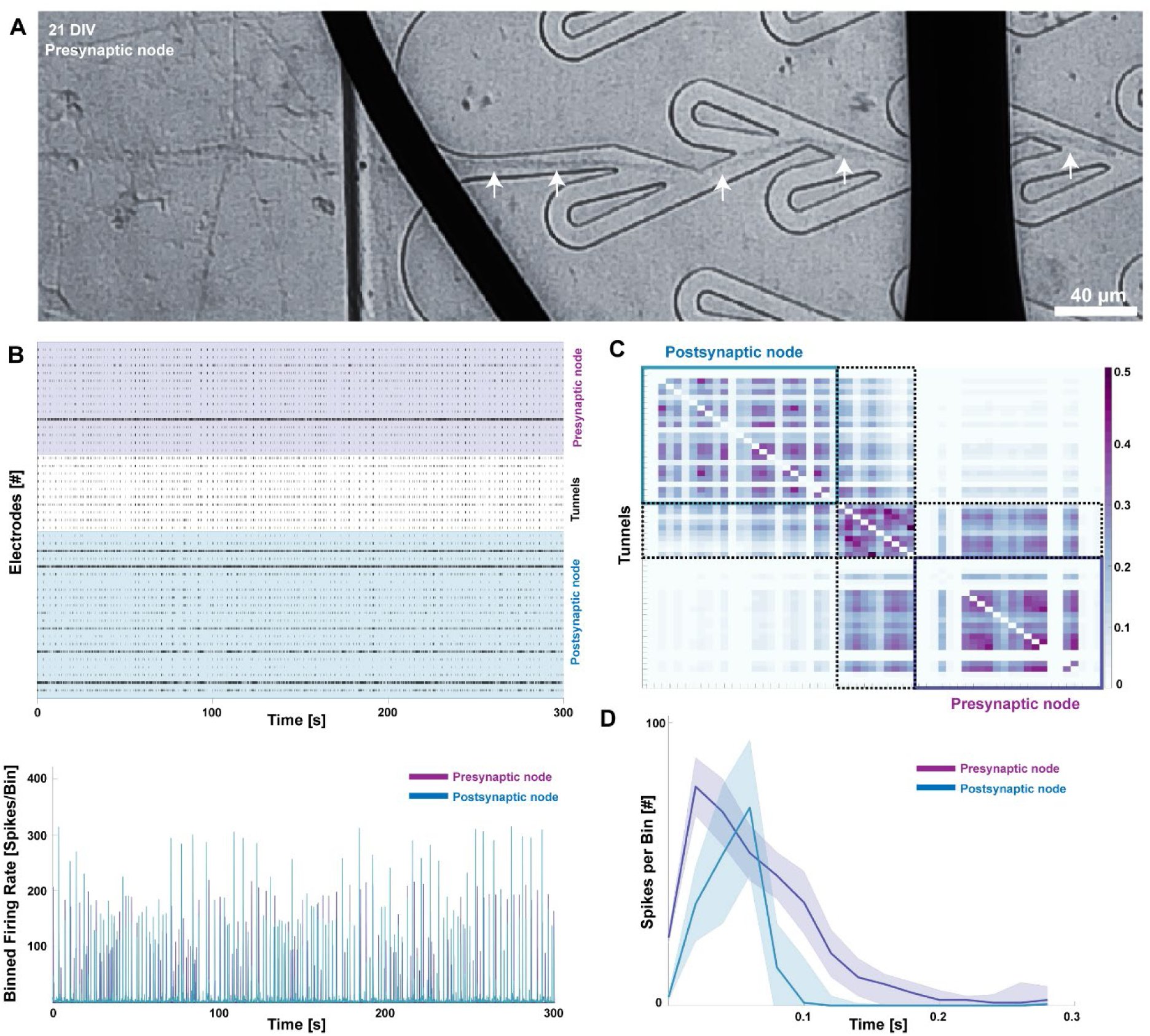
Developing structural organization with intra – and internodal functional connectivity of feedforward engineered neural networks. **(A)** Phase contrast images of a neural network in the presynaptic node with neurite projections going through the microtunnels on a mMEA at 21 DIV. **(B)** 300 s Raster plots of the recorded activity (top panel) and binned network activity of corresponding total firing rate (bottom panel) of one representative network at 26 DIV. **(C)** Connectivity matrix using mutual information to evaluate the correlations in the network activity within and across nodes. **(D)** Peristimulus time histogram of the pre- and postsynaptic response to electrical stimulations in the presynaptic node of a two-nodal network. The graphs show an initial response in the presynaptic node, followed by a delayed response in the postsynaptic node.

### 3.2. Perturbed networks express human mutated tau

At 28 DIV, we induced expression of human mutated tau protein in the presynaptic nodes of the six engineered networks. The AAV construct additionally encoded GFP, which allowed for easy visualization of transduction efficacy. Positive GFP expression was seen in the perturbed networks (**Figure 4B and Figure 5B**) which indicated effective AAV transduction. For additional verification of phosphorylated tau expression, we also labeled the networks for AT8 (Serine 202 and Threonine 205/ tau^202/205^) and Phospho-Tau (Threonine 217/tau^217^) proteins, hereon referred to as tau^202/205^ and tau^217^, respectively. Perturbed networks had significantly higher expression of both tau^202/205^ (**Figure 4B**) and tau^217^ (**Figure 5B**) in the cytosolic compartments (Rajbanshi et al., 2023) compared to controls, and a non-significant difference in expression levels in axon bundles. Primary exclusion immunolabeling for both tau^202/205^ and tau^217^ confirmed antigen specificity (**Figures 4C and 5C**). We also performed secondary antibody exclusion to assess the labeling specificity of the primary antibody and found no immunoexpression for either tau^202/205^ or tau^217^ in the networks (no data included).

**Figure 4.**
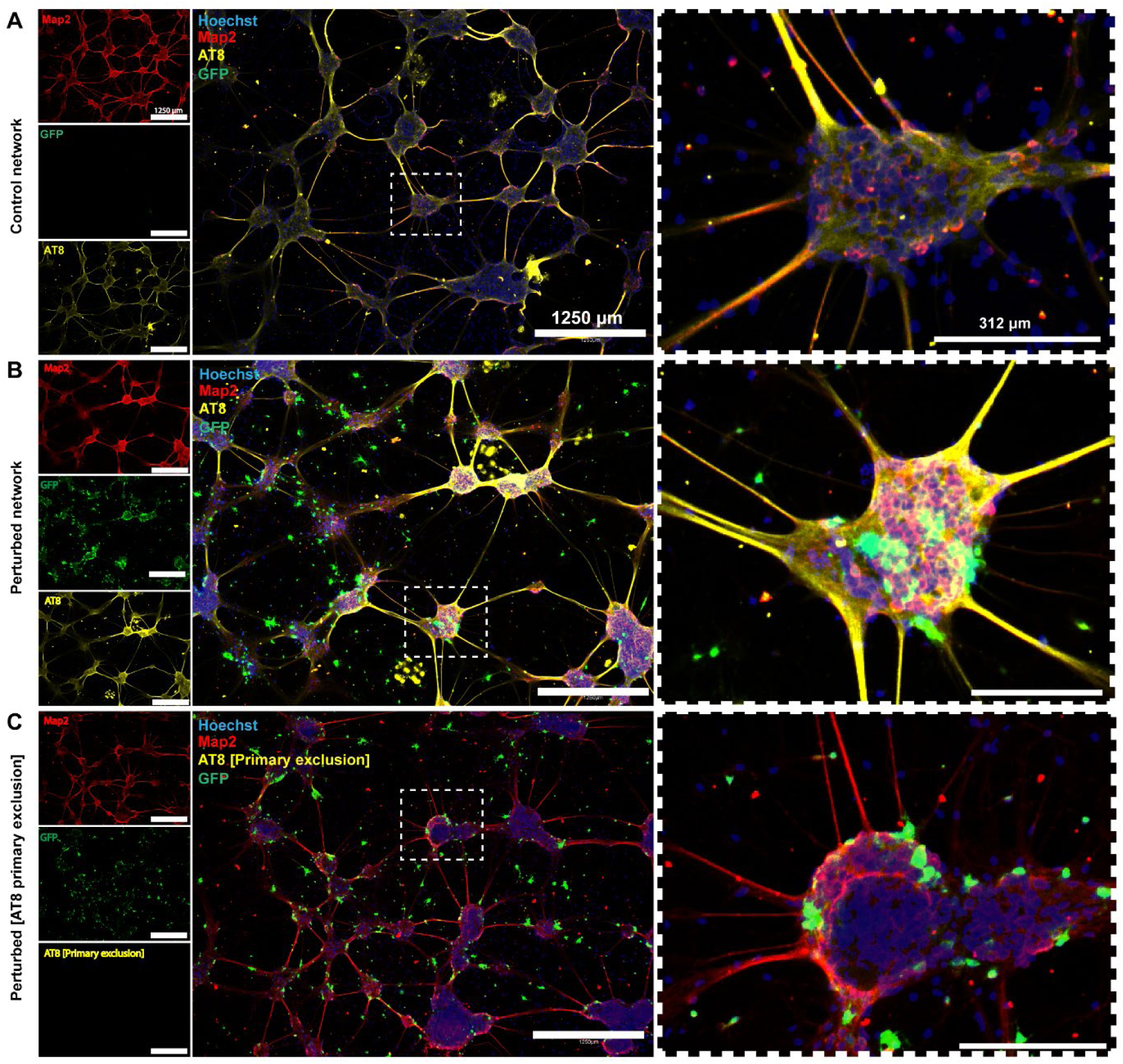
GFP expression exclusively in perturbed networks confirmed efficacy of viral transduction, further validated by AT8 expression. **(A)** Unperturbed control networks were not transduced with the AAV8-GFP-2A-P301L construct and thus did not express GFP. AT8 labeling was found primarily in axons. **(B)** Perturbed networks were positive for the GFP marker after transduction, with strong axonal and somato-dendritic AT8 expression. **(C)** Exclusion of the primary antibody AT8 to assess the specificity of antigen binding. Scale bar 1250 μm; (magnified area 312 μm).

**Figure 5.**
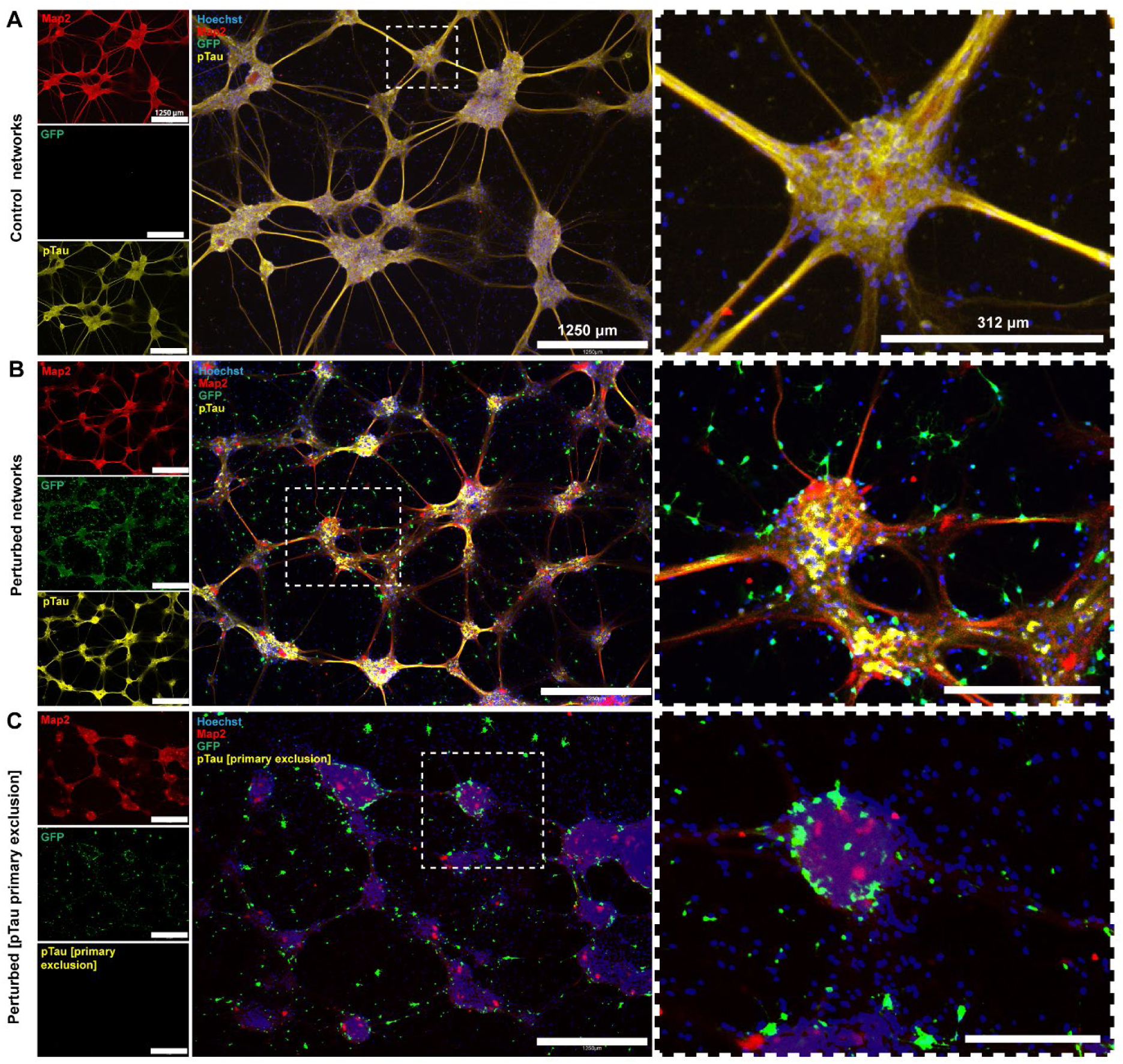
GFP expression exclusively in perturbed networks confirmed efficacy of viral transduction, further validated by pTau expression. **(A)** Unperturbed control networks were not transduced with the AAV8-GFP-2A-P301L construct and thus did not express GFP. pTau labeling was found in axons and cytosols. **(B)** Perturbed networks were positive for the GFP marker after transduction, with strong axonal and somato-dendritic pTau expression. **(C)** Primary exclusion of pTau to assess the specificity of antigen binding. Scale bar 1250 μm; (magnified area 312 μm).

**Figure 6.**
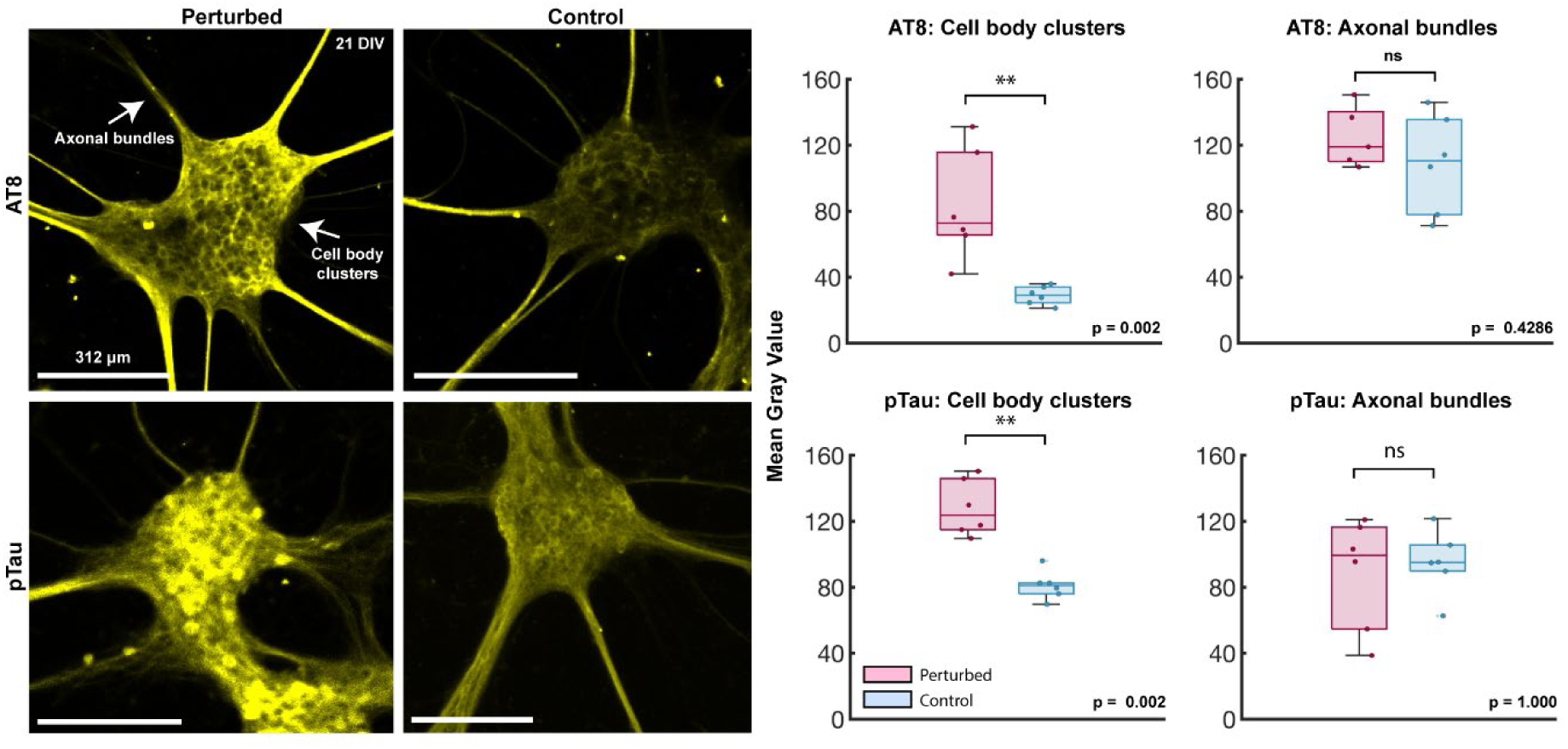
Quantification of fluorescent intensity of AT8 (tau^202/205^) and pTau (tau^217^) in perturbed and control unperturbed networks. Left: Representative images from networks with tau^202/205^ and tau^217^ labelling showing cell body clusters and axonal bundles. Right: Bar graphs showing the mean gray value of cell body clusters and axonal bundles of perturbed and unperturbed control networks. Cell bodies of perturbed networks show a significantly higher expression of both tau^202/205^ and tau^217^.

### 3.3. Progressive internodal axonal retraction observed after perturbation

Prior to induced perturbation, phase contrast imaging of neural networks at 21 DIV confirmed structural connections between the pre- and postsynaptic neural nodes (**Figure 7A**). By four days post perturbation, i.e., at 32 DIV, the neurites within the presynaptic node of the perturbed networks started retracting from the entry zone near the unidirectional microtunnels, and by 52 DIV all structural connections between the pre- and postsynaptic nodes had been lost entirely (**Figure 7A**). Extensive neurite retraction was also observed in the postsynaptic node of the perturbed networks by 52 DIV. In contrast, unperturbed control networks maintained robust neurite connections along the microtunnels throughout the experiment (**Figure 7B**).

**Figure 7.**
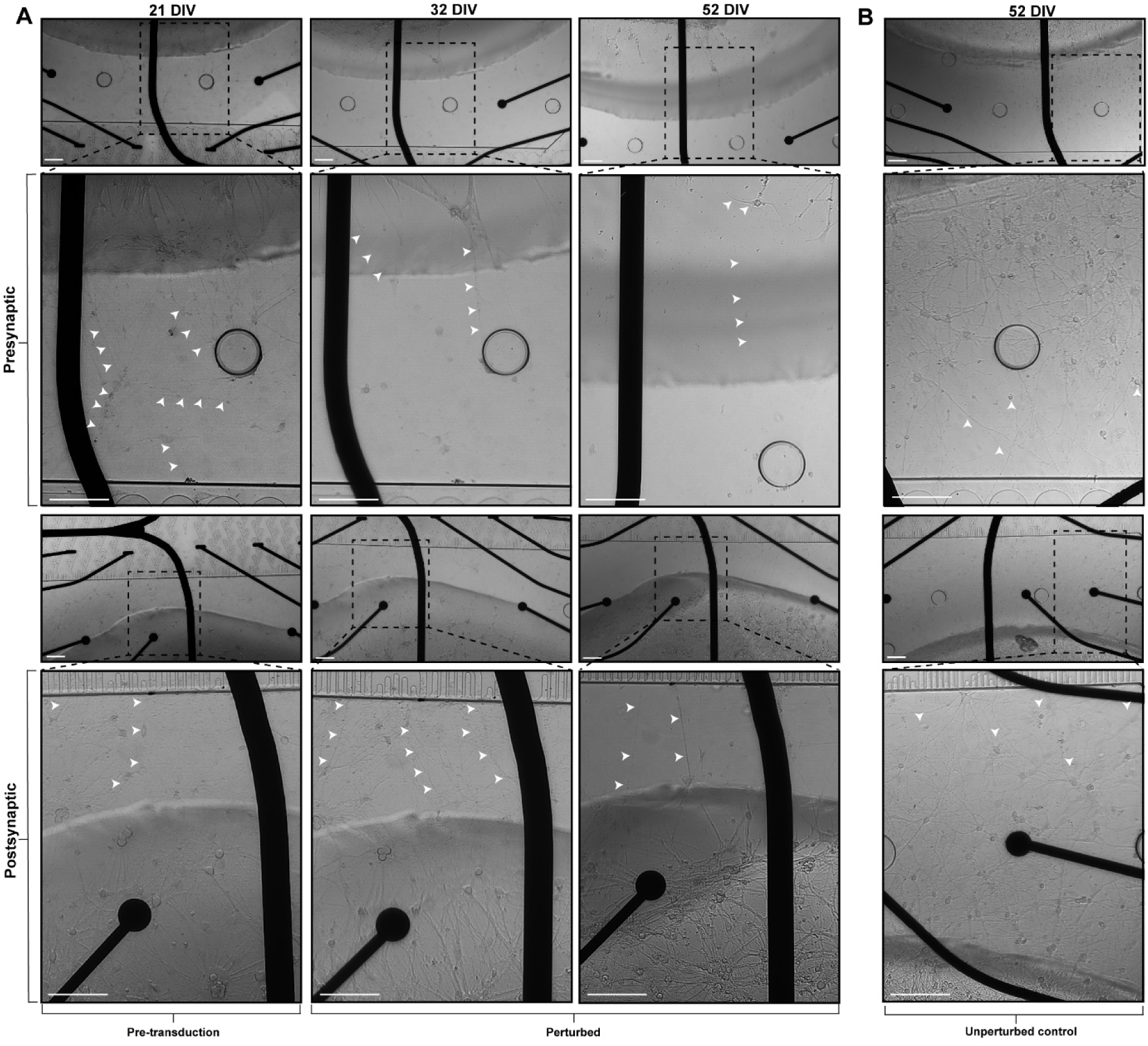
Progressive neurite retraction and active reorganization observed within networks following induced perturbation. **(A)** The leftmost column shows neurite extensions in the presynaptic node at the entry zone of the unidirectional microtunnels (top) and dense neurites in the postsynaptic node at the exit zone of the unidirectional microtunnels (bottom) at 21 DIV. The middle column is a snapshot of the same region in the network at 32 DIV showing retraction in the presynaptic node (top), but not in the postsynaptic node (bottom). The rightmost column shows extensive retraction in the presynaptic node (top) as well as in the postsynaptic node (bottom) at 52 DIV. **(B)** Presynaptic (top) and postsynaptic (bottom) nodes of an unperturbed control network depicting dense neurite connections towards the microtunnels at 52 DIV. Scale bars 100 μm.

### 3.4. Perturbed networks exhibit a decrease in overall network activity and an increase in network synchrony

Spontaneous network activity was recorded between 16 DIV and 47 DIV for both control unperturbed and perturbed networks. We captured network development from low activity towards more mature profiles exhibited as increased mean firing rate (**Figure 8A**) and increased mean burst rate (**Figure 8B**). The presynaptic nodes of unperturbed control networks exhibited a steady increase in electrical activity between 16 DIV and 31 DIV, consistent with previous studies by us and others capturing developing functional activity in cortical networks (Wagenaar et al., 2006; Weir et al., 2023). Both firing rate and burst rate increased between 33 DIV and 45 DIV in the presynaptic node but not in the postsynaptic node (**Figure 8A and B, respectively)**.

**Figure 8.**
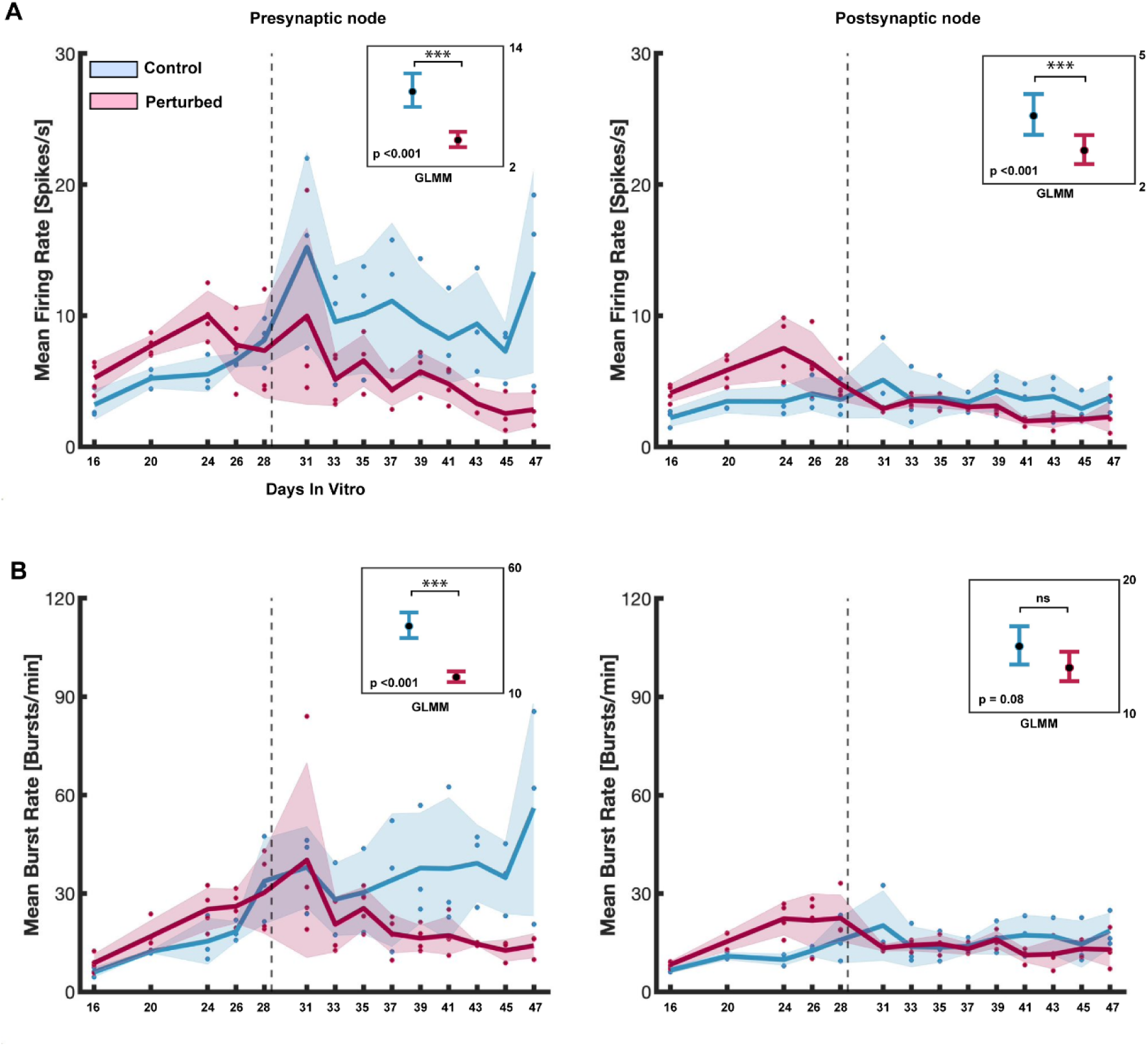
Electrophysiological recordings revealed progressively decreasing firing and burst rates in perturbed networks. **(A)** Mean firing rate (spikes/second) and **(B)** Mean burst rate (bursts/minute) in the pre- and postsynaptic nodes for unperturbed controls and perturbed networks. The solid lines denote the mean activity for the networks with the solid circles indicating individual data points. The shaded area denotes the standard error of the mean. The stippled line indicates the day of viral transduction in the perturbed networks (28 DIV). Plots showing the GLMM estimated group averages with 95% confidence intervals are depicted in the top right corner of the graphs.

Between 31 DIV and 47 DIV, both the pre- and postsynaptic nodes of perturbed networks exhibited significantly lower firing (p< 0.001) and burst (p< 0.001) rates compared to healthy controls (**Figure 8A**). There were no significant differences in mean burst rate in the postsynaptic node between healthy controls and perturbed networks (**Figure 8B**). The postsynaptic node of healthy controls had significantly longer bursts (p= 0.001) compared to perturbed networks (**Figure 9B**), however, there were no significant differences in the mean burst duration in the presynaptic nodes between perturbed and healthy controls between 31 DIV and 47 DIV (**Figure 9A**). Furthermore, the total network activity (pre- and postsynaptic node activity combined) revealed that between 28 DIV and 47 DIV, healthy controls had significantly higher mean network burst duration (p< 0.001) compared to perturbed networks (**Figure 9C**). Specifically, between 28 DIV and 33 DIV, unperturbed control networks exhibited a transient increase in mean network burst duration from 0.23 seconds to 0.26 seconds, however, there was a subsequent decrease between 33 DIV and 47 DIV (**Figure 9C**). For the perturbed networks, the total network activity for pre-and postsynaptic nodes combined showed that between 33 DIV and 39 DIV there was a transient increase in network burst duration from 0.17 seconds to 0.22 seconds, with a subsequent decrease between 39 DIV and 47 DIV similar to the healthy controls.

We also found that all networks had a general decrease in mean network burst size (analyzed as the percentage of active electrodes participating in network bursts), between 16 DIV and 28 DIV (**Figure 9D**). The total bursting activity for pre- and postsynaptic nodes combined showed that unperturbed control networks decreased in burst size from 85% to 45% network participation (**Figure 9D**) between 16 DIV and 28 DIV. Similarly, networks before perturbation had a decrease in burst size from 82% to 43% network participation (**Figure 9D**). Unperturbed control networks showed a relatively stable decrease in burst size between 28 DIV and 47 DIV. Contrarily, perturbed networks had significantly larger network bursts (p< 0.001) by 47 DIV (61% network participation) compared to control networks (45% network participation) (**Figure 9D**). Furthermore, when we examined network synchrony, measured as the coherence index, we observed a general decrease in both pre- and postsynaptic nodes of healthy controls until 31 DIV before it stabilized (**Figure 9E-F**). Both nodes of perturbed networks had progressively increased synchrony that was significantly higher (p< 0.001) than in control networks between 31 DIV and 47 DIV (**Figure 9E-F**).

**Figure 9.**
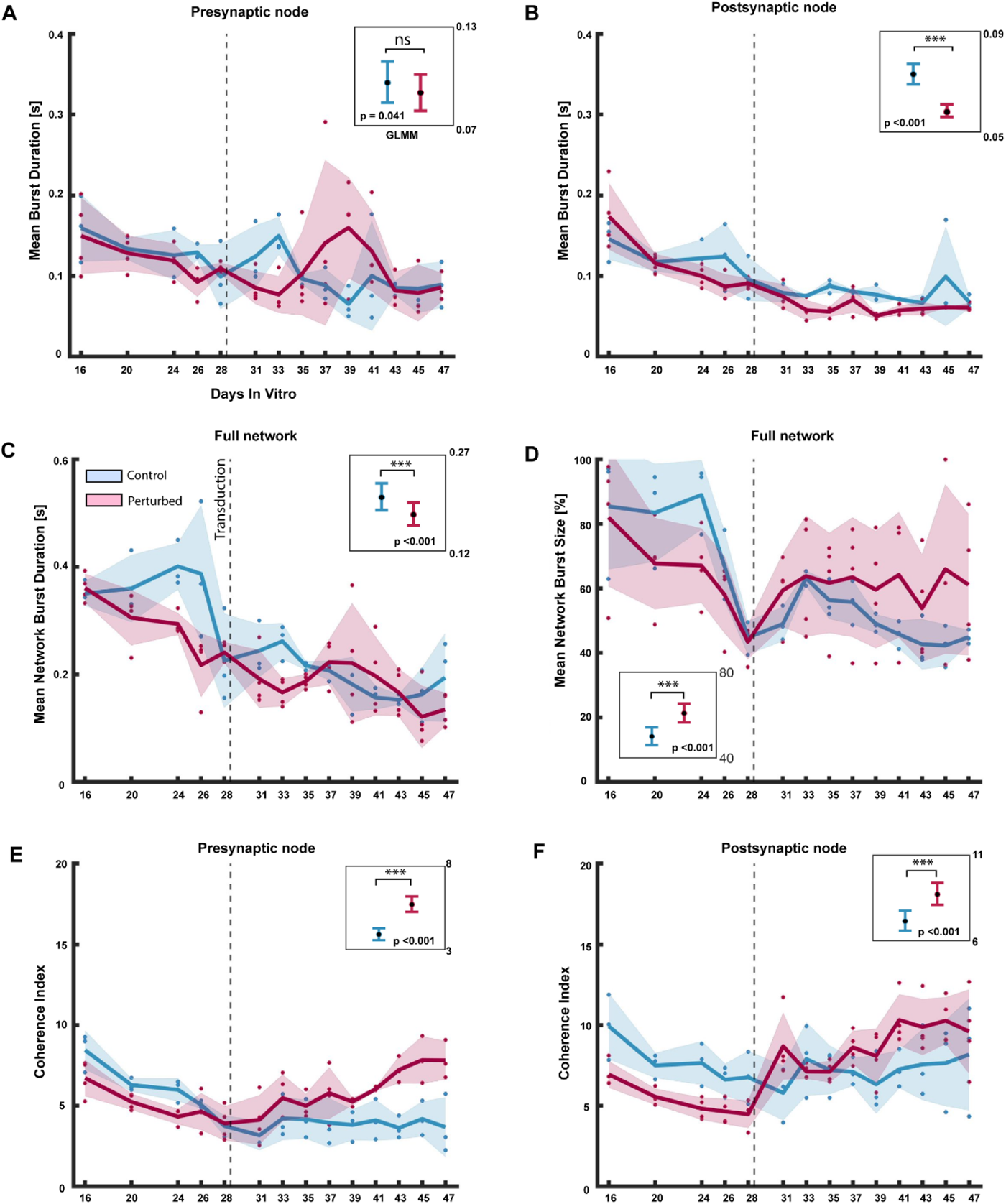
Electrophysiological recordings revealed a progressive increase in network burst size and synchrony in perturbed networks. (A) Mean burst duration (seconds) in the pre- and (B) postsynaptic nodes of control and perturbed networks. (C) Mean network burst duration in the pre- and postsynaptic nodes combined for control and perturbed networks. (D) Mean network burst size (percentage) in the pre- and postsynaptic nodes combined for control and perturbed networks. (E) Coherence Index (a measure of network synchrony) in the pre- and (F) postsynaptic nodes of control and perturbed networks. The solid line denotes the mean activity for the networks with solid circles indicating individual data points. The shaded area denotes the standard error of the mean. The stippled line indicates the day of transduction for the perturbed networks (28 DIV). Plots showing the GLMM estimated group averages with 95% confidence intervals are depicted in the top right corner of the graphs.

### 3.5. Induced perturbation results in reduced propagation of stimulation evoked activity between nodes

We also applied periodical electrical stimulations to the electrode with the highest firing rate in the presynaptic node to assess how an external presynaptic stimulus evoked a postsynaptic response. In each of the mMEAs, one electrode was stimulated for 1 minute at each recording session between 28 DIV and 47 DIV. We found that electrical stimulations within the presynaptic node of unperturbed control networks produced a response within the presynaptic node followed by a response in the postsynaptic node with an average delay time of 20 ms (31 DIV) and 40 ms (35 DIV) (**Figure 10A**). Unperturbed control networks also produced spike responses in the presynaptic node at 45 DIV and 47 DIV, although the tuning curves were of lower amplitudes than after previous stimulations. In addition, the spike responses in the postsynaptic node of control unperturbed networks at 45 DIV and 47 DIV were too low to allow for an evaluation of the delay time between the nodes. In contrast, stimulation in the presynaptic node of perturbed networks at 31 and 35 DIV resulted in a presynaptic spike response, with no spike response in the postsynaptic node (**Figure 10B**). There was no response in pre- nor postsynaptic nodes at 45 DIV and 47 DIV in the perturbed networks.

**Figure 10.**
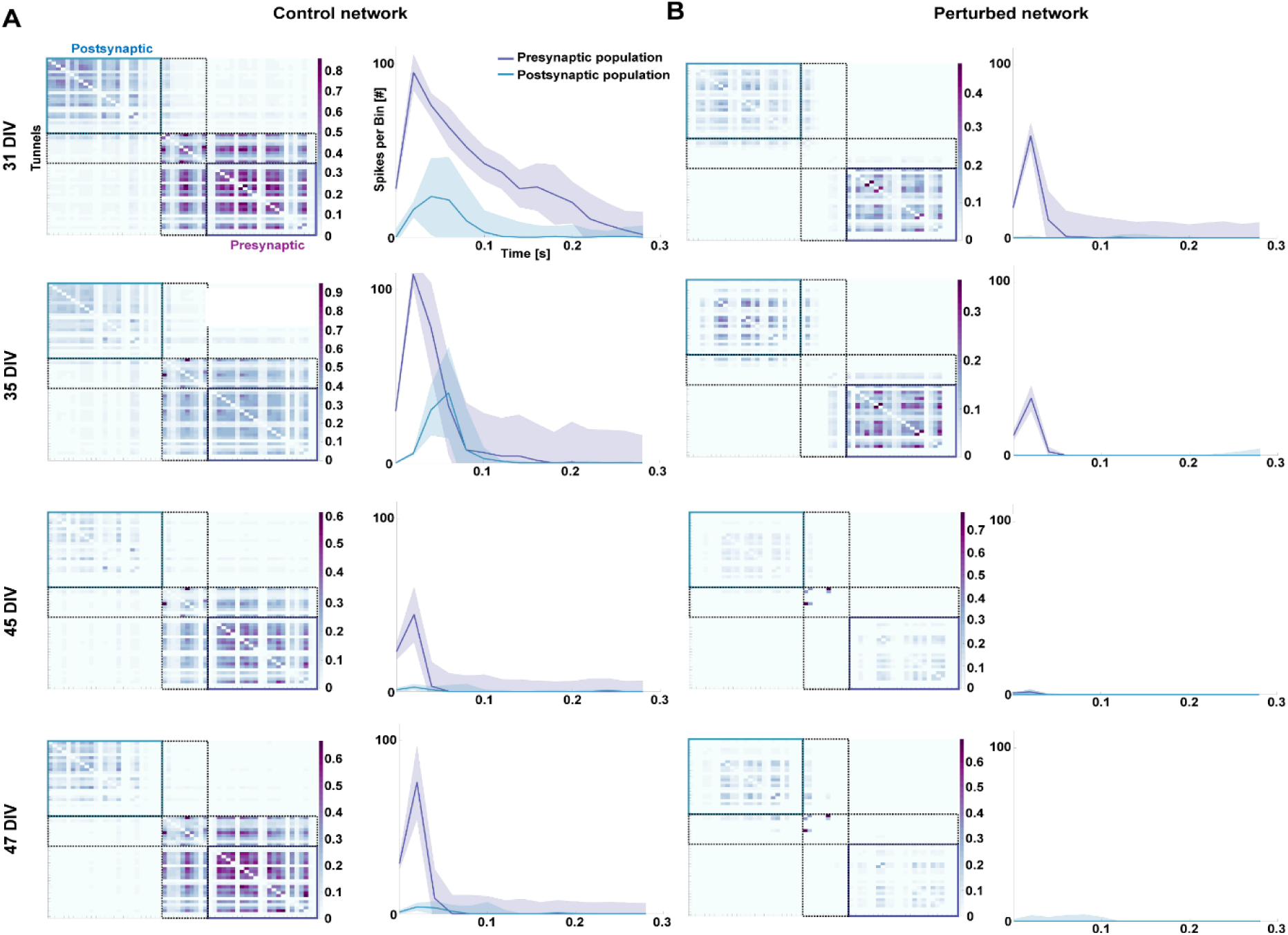
Induced perturbation resulted in progressive decline in presynaptic response to electrical stimulations between the pre- and postsynaptic nodes over time. Connectivity matrices for control (A) and perturbed (B) networks showing the mutual information between pairs of electrodes (left column) and peristimulus time histograms of pre- and postsynaptic responses to electrical stimulations (right column) at 31, 35, 45 and 47 DIV. The curves show an initial response in the presynaptic node, followed by a delayed response in the postsynaptic node for control networks and no postsynaptic response for perturbed networks. The shaded area denotes the standard error of the mean of 60 repeated stimulations.

## 4. Discussion

Engineered neural networks *in vitro* self-organize into complex topologies and produce complex profiles of activity patterns ranging from individual spikes produced by single neurons to high frequency network bursts generated by neural assemblies (Chiappalone et al., 2006; van de Wijdeven et al., 2018; Weir et al., 2023). The development of complex network activity can be attributed to the structural properties of the network, as function tends to co-evolve with structure (Kapucu et al., 2017). Activity typically evolves with the maturity of neurons, and of excitatory and inhibitory synapses. Our engineered neural networks expressed both the excitatory neuronal marker CaMKIIa, and the inhibitory neuronal marker GAD65/67 in conjunction with the neuronal marker MAP2 by 21 DIV, strongly indicating the capacity for excitatory – inhibitory synaptic transmission within the maturing networks. This was further verified through recordings of electrophysiological activity, which showed that in healthy conditions, networks developed highly dynamic and complex age-dependent firing activity and network bursts over time, in line with previous studies by us and others (Chiappalone et al., 2006; Chiappalone et al., 2007; Fiskum et al., 2021; van de Wijdeven et al., 2018; Weir et al., 2023).

The structural organization of neural networks is a crucial determining factor for the emergence of complex network dynamics and information processing. In our engineered feedforward networks, we showed that neurites extended from the presynaptic nodes into the microtunnels towards the postsynaptic nodes, thus establishing structural connections. Furthermore, electrical stimulation at 26 DIV confirmed that these networks were functionally interconnected, as evoked activity in the presynaptic node could propagate and elicit a response in the postsynaptic node, in line with previous findings (Fair et al., 2009; Ma et al., 2018; Winter-Hjelm N, 2023). These results confirm recapitulation of a feedforward network *in vitro*, where information flows in a unidirectional manner, propagating sequentially from input nodes to output nodes (Barzegaran et al., 2022; Markov et al., 2014). Such feedforward hierarchical organizations are found in many parts of the brain (Markov et al., 2014; Siegle et al., 2021), and facilitate fast, efficient information processing between pre- and postsynaptic neuronal assemblies. The ability of the network to spontaneously transmit information between the nodes is an essential factor in determining its functional capacity since this enables the integration of signals that support coordinated, complex information processing (Fauth et al., 2019; Fukushima et al., 2018; Senden et al., 2018). As such, our results confirmed the intended structural and functional organization of engineered feedforward cortical networks, with the presynaptic node providing input to the postsynaptic node.

These engineered feedforward networks also enabled us to selectively induce a perturbation, via AAV mediated expression of human mutated tau in the presynaptic node, and to monitor resulting effects within and between nodes. By utilizing this approach, we can effectively recapitulate and monitor a pathological process at the micro- and mesoscale. Such a process not only induces structural and functional changes in the immediately affected node, but also disrupts both structure and function in the postsynaptic node (Kuhl, 2019; Valderhaug et al., 2021). These findings are highly relevant for advanced modelling of evolving pathological processes in neurodegenerative diseases, such as AD, where an association between the progression of tau pathology and altered transsynaptic activity has been demonstrated (Liu et al., 2012). Such transsynaptic spread of tau pathology is attributed to the robust connection loops that exist between the putative origin points of tau pathology in the lateral entorhinal cortex layer II (Braak et al., 1991) and feedforward axonal projections to hippocampal subregions (Liu et al., 2012; Stepan et al., 2015; Witter, 2007; Witter et al., 2017). This feedforward transsynaptic spread of hyperphosphorylated tau may contribute to structural impairments that disrupt the normal functioning of neural networks. In our study, we found prominently different tau^217^ and tau^202/205^ expression in the cytosol of neurons in perturbed networks compared to the controls, signifying differential consequences of tau phosphorylation for network function. Phosphorylation of tau^202/205^ is found in maturing brain and is associated with neural development (Goedert et al., 1995), while tau^217^ phosphorylation is found to be associated with the normal development of postsynaptic sites (Rajbanshi et al., 2023). This means that some phosphorylation of tau at these sites is to be expected in unperturbed control networks. Significantly, studies have shown that the phosphorylation of tau at a single site does not preferentially induce neurotoxic effects (Steinhilb et al., 2007), though recent findings reported a correlation between high tau^217^ expression and AD pathology progression (Rajbanshi et al., 2023) associated with synaptic decline. Our finding of increased expression of both tau^202/205^ and tau^217^ in the cytosol of neurons within the perturbed networks suggests that the neurotoxic effects of tau depend on a combined high phosphorylation pattern at multiple sites in axons and/or the cytosol. Furthermore, the decreased activity in the postsynaptic node of perturbed networks indicated spread of pathology between the nodes, possibly attributed to the increased phosphorylation of tau in the presynaptic neurons.

Other studies have found that the severity of neuronal loss and atrophy of cortical structures as a result of tau pathology tend to positively correlate with the severity of functional network decline (Adamec et al., 2002; Rascovsky et al., 2005). The present study revealed such dynamic structural and functional changes in perturbed networks, while highlighting the complex interrelationship between network structure and function in tau pathology. Between 32 DIV and 52 DIV, we observed progressive neurite retraction from the entry zone near the microtunnels in the presynaptic nodes in the perturbed networks. We also noticed that retraction from the exit zone near the postsynaptic nodes also occurred within weeks of presynaptic retraction. This was not observed in the unperturbed control networks, which maintained a dense neurite network in the zones near the microtunnels in both nodes. The significance of these changes in perturbed networks lies in their potential to adversely affect the network’s ability to transmit and integrate information. Prolonged expression of mutant P301L tau exacerbates axonal destabilization (Biswas et al., 2018; Qiang et al., 2018) and impairment of presynaptic terminals (Hunsberger et al., 2021). Furthermore, both maintenance of presynaptic integrity and synaptic plasticity depend on active anterograde and retrograde axonal transport systems (Cai et al., 2011). As such, impairment of tau in axons can affect such processes (Lacovich et al., 2017) and severely disrupt synaptic connectivity between pre- and postsynaptic nodes in perturbed networks. The observed subsequent neurite retraction in the postsynaptic node of the perturbed networks suggested that the loss of presynaptic input may have triggered reorganization within postsynaptic nodes. These observations thus indicated a dynamic process of structural and functional reconfiguration across the entire feedforward network in response to perturbation of the presynaptic node.

The prominent structural changes observed following induced perturbation occurred concomitantly with changes in the electrophysiological profile of the perturbed networks between 28 DIV and 47 DIV. In healthy conditions, gradually increasing firing and burst patterns are crucial for the establishment and maintenance of functional synapses, and the elaboration of the network topology into hierarchical processing. Bursts also contribute to the formation and refinement of neural circuits especially during early network formation where they facilitate the integration of immature neurons into the maturing network (Wagenaar et al., 2006). Following the induced expression of human mutated tau, perturbed networks exhibited a steady decline in both firing rate and burst rate in comparison to control unperturbed networks, which continued to display a steady increase over time. Furthermore, these differences in firing rate following perturbation were found to be significant between control unperturbed and perturbed networks in both pre- and postsynaptic nodes. This suggested that perturbed networks became less electrophysiologically active as they underwent structural reorganization, including neurite retraction. The observed decrease in firing and bursting activity also align with *in vivo* findings showing that neurons in a mouse model with the Tau-P301 mutation gradually became more hypoactive (Busche et al., 2019). However, Busche et al., (Busche, 2019) also suggested that the disruption in network activity occurred before any prominent structural tau abnormalities were observed *in vivo*. We have however shown that the decline in general network activity correlated strongly with the progressive loss of synaptic connectivity and pre-and postsynaptic neurite reorganization. These changes are highly challenging to detect and correlate *in vivo*.

Another interesting result in our study was the significant increase in network burst size and synchrony observed in the perturbed networks between 28 DIV and 47 DIV. Network bursts, which are coordinated patterns of neuronal activity exhibited by multiple interconnected neurons within the network, are ubiquitous for normal network function (Weir et al., 2023). Coordinated neuronal activity also leads to network synchrony (Salinas et al., 2001), and is thus important for information processing, coding, and synaptic integration of distributed signals (Gansel, 2022). Synchrony may also promote activity-dependent establishment of synaptic connections via spike-timing dependent plasticity (STDP) (Anisimova et al., 2022) to support network function. In a recent study by our group, we found that networks that were perturbed by selective silencing of excitatory synaptic transmission also demonstrated increased synchrony during network recovery (Weir et al., 2023). Such behavior may thus be crucial for the network’s ability to restore its functional and structural organization within specific time windows after a perturbation. On the other hand, increased synchrony may also have adverse effects, such as facilitating the spread of perturbations through axons and synapses to affect the entire interconnected network (Uhlhaas et al., 2006), as has been found in AD pathology (Liu et al., 2012; Wang et al., 2017). Furthermore, excessive network bursts and synchrony have been implicated in various neurological disorders, including epilepsy, where they signify a disruption in normal network physiology, i.e., impaired excitatory-inhibitory dynamics (Kudela et al., 2003; Wu et al., 2015). Therefore, while a degree of network synchrony is necessary for normal functioning, too much can be problematic. This raises a crucial question regarding whether the observed synchrony in the perturbed networks in our study may represent an adaptive or maladaptive network response to induced perturbation.

Interestingly, synchrony increased concurrently with neurite retraction at 32 DIV, while firing and burst rate declined. We found that, following perturbation, networks began exhibiting fewer, yet larger synchronized bursts, which can be interpreted as a homeostatic compensatory response to maintain network activity as the overall firing and burst rates declined. In response to low network activity levels, homeostatic scaling, which occurs gradually and over several hours to days, can increase overall input to counteract hypoactivity (Chowdhury et al., 2018; Turrigiano, 2008). This may explain the observation of increasing synchrony between 31 DIV and 47 DIV. However, increased synchrony could also signify pathophysiological changes in the underlying network since it occurred concomitant with the evolution of induced pathology. *In vivo,* induced perturbation caused by hyperphosphorylated tau can disrupt the structural and functional integrity of the affected network, and subsequently result in increased inflammation leading to apoptotic or necrotic cell death (Dong et al., 2022; Thal et al., 2022). Furthermore, it has been suggested that the presence of diverse connections and pathways within a neural network can provide alternative routes for information flow to reduce the reliance on a single synchronized pathway (Kirst et al., 2016), thus acting as a gatekeeping mechanism to prevent excessive synchrony and enhancing overall network robustness. In our study, as induced perturbation led to progressive structural and functional disruption in the network, it is likely that information flow within the network might have been hindered, as there were insufficient alternative routes to effectively distribute activity, ultimately leading to excessive synchrony within the individual nodes. This is further supported by observations in unperturbed networks between 31 DIV and 47 DIV. During this time, control networks exhibited significantly higher mean firing and burst rates without showing excessive synchrony. These networks also maintained the dense neurite architectures between the nodes. This effectively suggests that the unperturbed control networks possessed the ability to maintain their activity within a dynamic range to prevent excessive network wide activation, an ability which the perturbed networks appeared to gradually lose. Based on these findings, increased network synchrony after induced perturbation might be associated with the deterioration of overall network function in response to the perturbation, rather than serving as an adaptive purpose.

Lastly, to further investigate whether perturbation affected structural and functional connectivity between the nodes, we applied periodic electrical stimulation to one electrode within the presynaptic node and assessed the postsynaptic response. We found that although unperturbed networks exhibited a consistent presynaptic spike response to electrical stimulation, there was a gradual decline in the postsynaptic response over time. This could be due to activity dependent long-term synaptic changes in the vicinity of the stimulating electrode, such as a reduction in synapse number or downscaling of synaptic receptors on neurons, as previously reported (Collingridge et al., 2010). Such activity dependent structural changes would likely reduce the amplitude of the presynaptic response, thus reducing the strength of the propagating signal. For perturbed networks, we found that there was no response to presynaptic stimulation in the postsynaptic node by 31 DIV, with no response in the presynaptic node at 45 DIV and 47 DIV. This outcome may be attributed to the extensive neurite retraction in the presynaptic node indicating the severance of connectivity between nodes, a possible progressive neuron loss in the network or even or structural reorganization of any remaining neurons to areas outside the vicinity of the stimulating electrode.

## 5. Conclusions and future directions

This study utilized engineered two-nodal feedforward neural networks with controllable afferent-efferent connections to longitudinally monitor and assess dynamic structural and functional behaviors in healthy conditions and in response to induced perturbation via the expression of human mutated tau. The findings revealed that prior to perturbation, both unperturbed and perturbed networks followed similar developmental trajectory consistent with relevant literature. The effects of the induced perturbation were evident within one week, with perturbed networks exhibiting a significant decrease in firing rate, burst rate and total number of bursts in contrast to the increases observed in control unperturbed networks. These changes are consistent with the known adverse effects of tau hyperphosphorylation. Over time, while control networks showed a decline in burst size and synchrony, perturbed networks demonstrated significant increases in both metrics, suggesting maladaptive synchrony responses linked to neurite retraction and reduced overall network activity. Importantly, these changes were not seen in healthy unperturbed controls, confirming that they were due to the induced perturbation rather than endogenous tau expression.

The significance of this study lies in its successful monitoring of ongoing structural and functional changes at the micro and mesoscale network level, and providing access to processes that are otherwise challenging to study *in vivo*. The findings also underscore the platform’s efficacy in investigating perturbation-induced changes, providing insights into tau-associated pathology and offers significant potential for examining molecular-level effects, such as mitochondrial changes, which were not within the scope of the current study but are crucial for future research. Another valuable enhancement for future study will also involve introducing a control virus featuring a wild-type tau with the same structure as the experimental virus to enable a more thorough comparison of the effects induced by mutated tau. While the current control unperturbed networks served as a sufficient baseline for evaluating structural and functional changes, this addition would strengthen the study’s robustness.

Overall, our current results provide significant new insights into the dynamic structural and functional reconfigurations in neural networks caused by evolving tau pathology. These findings have profound implications for future research, enhancing the ability to track and analyze intricate neural processes over time, and paving the way for advancements in understanding and treating progressive neuropathological processes.

## Author Contributions

The author contribution follows the CRediT system. JSW, KSH: Conceptualization, Methodology, Investigation (Cell experiments; Protocol development and optimization; AAV investigations, Immunocytochemistry and Electrophysiological recordings), Writing – Original Draft, Review and Editing, Illustration. NWH: Methodology, Software, Investigation (chip design & manufacturing, electrophysiology, formal analysis), Writing – Review and Editing. AS, IS: Conceptualization, Methodology, Writing – Review & Editing, Funding acquisition, Supervision.

## Acknowledgements

The authors would like to thank Dr. Christiana Bjørkli for the technical support in developing the transduction protocol and gifting us the AAV8-GFP-2a-P301Ltau-virus, originally designed, and produced by Dr. Rajeevkumar Nair Raveendran at the Viral Vector Core Facility, Kavli Institute for Systems Neuroscience. Prof. Michela Chiappalone and Prof. Sergio Martinoia, University of Genova for generously providing the scripts for the Precise Timing Spike Detection algorithm and the logISI burst detection. Prof. Menno P. Witter and Dr. Asgeir Kobro-Flatmoen, Kavli Institute for Systems Neuroscience for kindly reading and proving feedback.

## Data availability statement

The raw data that support the findings of this study will be made available by the authors upon request.

